# Spotlight on islands: on the origin and diversification of a new lineage of the Italian wall lizard *Podarcis siculus* in the western Pontine Islands

**DOI:** 10.1101/293985

**Authors:** Senczuk Gabriele, Havenstein Katja, Milana Valentina, Ripa Chiara, De Simone Emanuela, Tiedemann Ralph, Castiglia Riccardo

**Affiliations:** Department of Biology and Biotechnology ‘Charles Darwin’, University of Rome La Sapienza, Italy; Unit of Evolutionary Biology/Systematic Zoology, Institute of Biochemistry and Biology, University of Potsdam, Karl-Liebknecht-Strasse 24–25, Haus 26, 14476 Potsdam, Germany

## Abstract

Groups of proximate continental islands may conceal more tangled phylogeographic patterns than oceanic archipelagos as a consequence of repeated sea level changes, which allow populations to experience gene flow during periods of low sea level stands and isolation by vicariant mechanisms during periods of high sea level stands. Here, we describe for the first time an ancient and diverging lineage of the Italian wall lizard *Podarcis siculus* from the western Pontine Islands. We used nuclear and mitochondrial DNA sequences of 156 individuals with the aim of unraveling their phylogenetic position, while microsatellite loci were used to compare several a priori insular biogeographic models of migration with empirical data. Our results suggest that the western Pontine populations colonized the islands early during their Pliocene volcanic formation, while populations from the eastern Pontine Islands seem to have been introduced recently. The inter-island genetic makeup indicates an important role of historical migration, probably due to glacial land bridges connecting islands followed by a recent vicariant mechanism of isolation. Moreover, the most supported migration model predicted higher gene flow among islands sharing a longitudinal arrangement. Considering the threatened status of small insular endemic populations, we suggest this new evolutionarily independent unit be given priority in conservation efforts.

## Introduction

Islands epitomize a suite of simplified conditions in spatially bounded areas, which make harbored populations an ideal model to investigate several evolutionary topics^1–3^. For example, oceanic islands contributed a lot to our empirical understanding of a variety of microevolutionary processes that drive diversification and eventually speciation^4–7^. In such a context, a pioneer species that colonizes an oceanic archipelago through oversea dispersal is expected to mold genetic structure mainly in function of island size, physiography, proximity among islands and dispersal capabilities. Conversely, islands situated on continental shelves have often been reconnected with the mainland after sea level drops during glacial phases^8^. This scenario creates allopatric conditions that change through time, enabling both vicariant and dispersal mechanisms to act on genetic diversity and divergence^9,10^. In addition, groups of small geographically proximate islands may comprise an even more complex phylogeographic setting compared to single continental islands because nearby islands may experience repeated cycles of isolation (during sea level transgression) and connection (during periods of low sea level stands). Indeed, a complex geological setting dominated by considerable changes to island size might lay the foundation for a variety of different genetic outcomes as a result of changing demographic processes and recurrent periods of fragmentation and gene flow^11–13^.

With almost 5000 islands, the Mediterranean Basin provides an attractive geological history which helps to explain the insular biodiversity observed today^14^. During the Pleistocene, the basin was characterized by cyclic sea level drops, which caused the formation of transient land bridges connecting nearby islands to each other, as well as islands and the mainland. The Mediterranean insular biota contains numerous taxa formed in response to these geological fluctuations^15–17^.

Almost half of the 23 recognized species of *Podarcis* lizards are endemic to Mediterranean islands and they represent an ideal model to explore the evolution of genetic diversity on islands. Here, we focus on the Italian wall lizard *Podarcis siculus* which, aside from inhabiting the entire Italian Peninsula, is also widespread across the greater islands of Sicily, Corsica, and Sardinia and across all the main Italian archipelagos and smaller islands (Fig. 1a). The phylogeography of this species across the Italian Peninsula is very complex^18^ and a recent study^19^ revealed that long term processes, giving rise to a mosaic of allopatric lineages, combined with more recent demographic dynamics have driven the evolutionary history of this species. Here, we examine insular genetic variation of *P. siculus* from the Pontine Archipelago using mitochondrial sequences (mtDNA), nuclear sequences (nuDNA), and microsatellite loci. The Pontine Archipelago is formed by two groups of islands. The western group, which lies 32 km from the Tyrrhenian Coast, includes the oldest island of Ponza and the nearby Gavi Islet, Zannone Island, and Palmarola Island. These islands are separated from the eastern Pontine Archipelago by 40 km of open sea; this eastern group of islands includes Ventotene and the Santo Stefano islands, located 50 km away from the mainland (Fig. 1b). On the basis of morphological traits, different subspecies have been described within this archipelago: *Podarcis s. latastei* (Bedriaga, 1879) on Ponza Island*, Podarcis s. patrizii* (Lanza, 1952) on Zannone Island*, Podarcis s. lanzai* (Mertens, 1952) on Gavi Islet, and *Podarcis s. palmarolae* (Mertens, 1967) on Palmarola Island.

**Figure 1.**
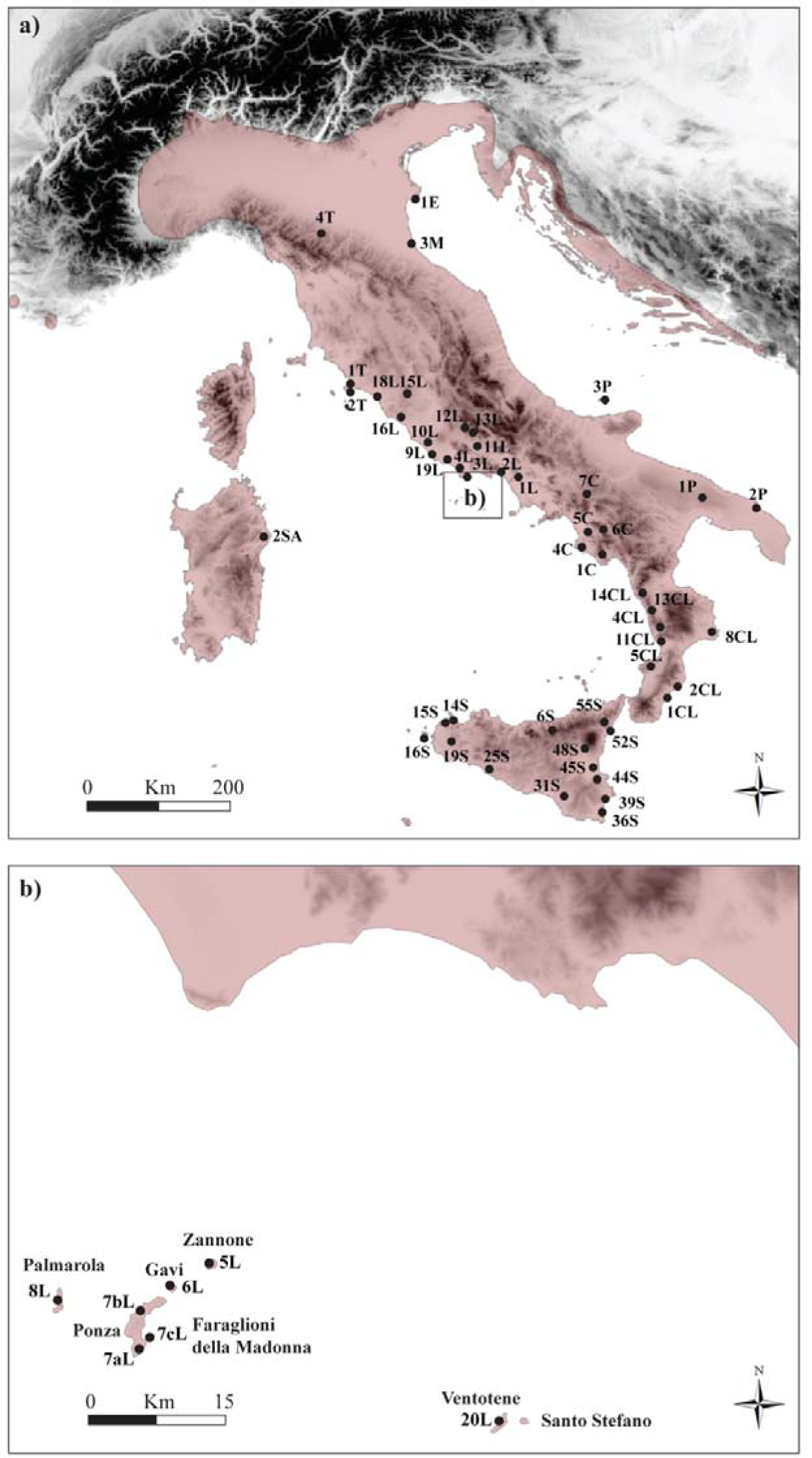
**(a)** Map of the study area showing all sampling sites on the Italian Peninsula with the geographic distribution range of *Podarcis siculus* highlighted in red. **(b)** Map of the Pontine Archipelago with relative sampling locations.

This particular insular context provides ample opportunity to investigate the origin and evolution of intraspecific genetic diversity within a Mediterranean archipelago for at least several crucial factors. First, the volcanic origin of the islands has been thoroughly investigated and relatively precise ages for their emergence have been reported. Presumably, there never experienced any stable contact with the mainland^20,21^. Second, the surrounding shallow waters indicate that the islands have been subjected to repeated episodes of fragmentation during high sea level stands as well as reunification when sea levels were lowest, during glacial phases. Finally, the western Pontine Islands have an attractive spatial arrangement. The bigger island of Ponza is located in the middle of the archipelago and the smaller Palmarola and Zannone islands show different longitudinal settings. All these features make this island system an ideal model to test different hypotheses of island biogeography, specifically regarding their compatibility with the observed patterns of genetic divergence and differentiation in this Mediterranean archipelago.

In this study we aim to (i) unveil the evolutionary history of *Podarcis siculus* insular populations, and (ii) assess the influence of different scenarios of inter-island genetic diversification and gene flow by comparing different theoretical models of island biogeography with genetic empirical data.

## Results

### Phylogenetic reconstruction and time of divergence

In the concatenated mtDNA alignment of 1701 bp (*nd4* = 766 bp and *cytb* = 935 bp;156 sequences; accession numbers are reported in Supplementary Table S4), we found 100 haplotypes with 368 polymorphic sites and an overall mean distance of 0.062 between the main clades. Uncorrected pairwise *p*-distances are reported in Supplementary Table S5.

The phylogenetic inference returned the same topology under the two Bayesian approaches applied, unraveled the presence of eight clades (Fig. 2a and Supplementary Fig. S1), seven of which already described in previous works^18,19^. However, a new diverging lineage (clade P in Fig. 2a, b) was discovered to which all individuals from the western Pontine Islands belong. The phylogenetic reconstruction indicated an early split separating the S, C1, and C2 clades from all the other clades (A1, A2, T1, T2, and P). Subsequently, a second cladogenetic event involved the separation of the Pontine clade P.

**Figure 2.**
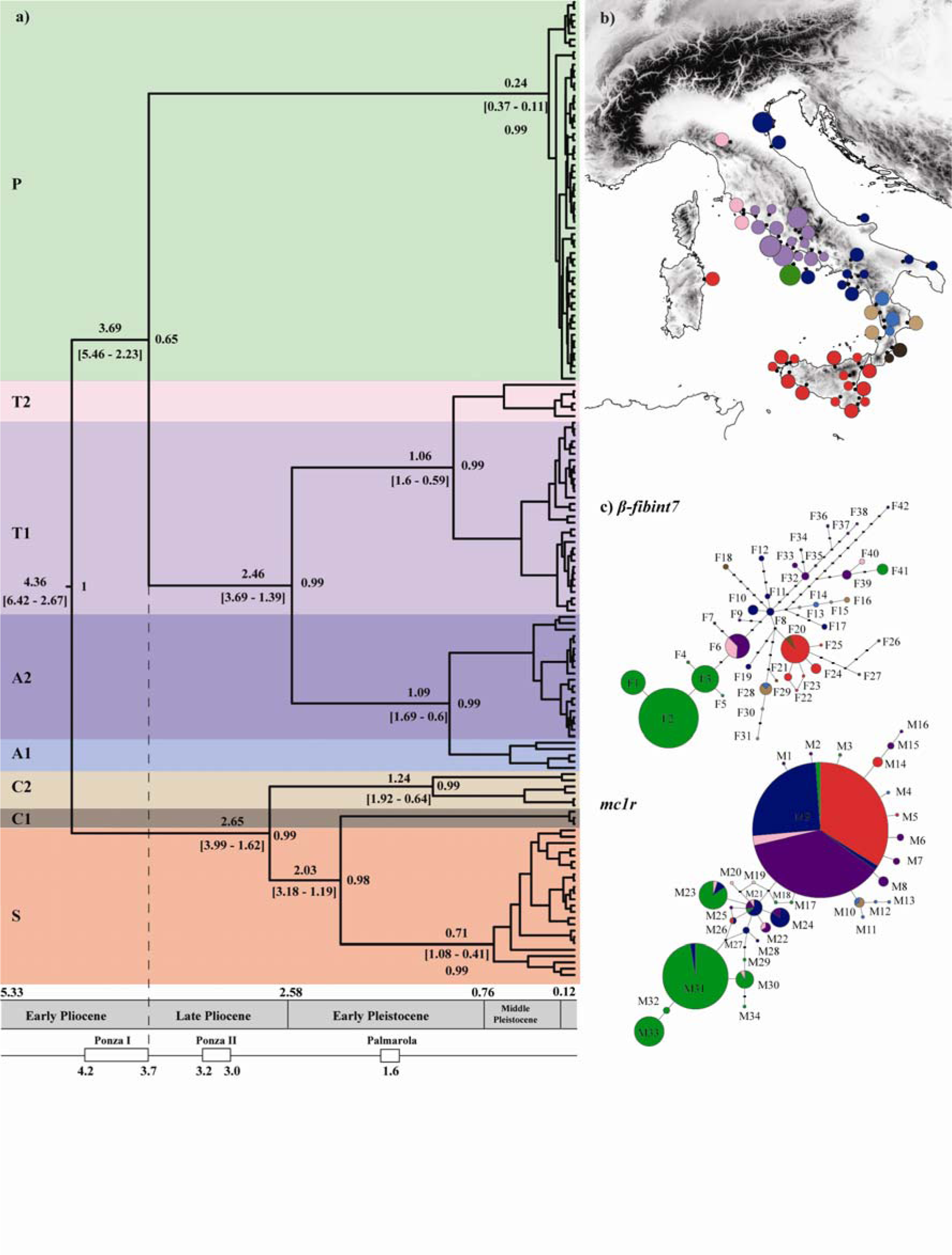
**(a)** Chronogram based on mtDNA performed with BEAST. At each node, the posterior probabilities and the coalescence time with their relative 95% high posterior densities (HPD) are shown. Chronology of the geological epochs with the two principal volcanic episodes that formed the western Pontine Islands shown below. **(b)** Geographic distribution of the mtDNA clades at each sampled locality. **(c)** Statistical parsimony networks of the two nuclear gene fragments; circle sizes are proportional to allele frequencies and filled rectangles representing one mutation step. Alleles are colored based on the mtDNA clade assignment of the respective specimen.

The divergence time analysis carried out using BEAST showed an effective sample size for all parameters over 200, indicating good chain mixing and adequate posterior estimates. The coefficient of variation in the average mutation rate estimates suggests a substantial heterogeneity among clades, supporting a non-clock-like model for the phylogeny (*ucld.stdev* parameter was 0.33 for *cytb* and 0.47 for *nd4*). According to our divergence time estimate, the Pontine clade (P) separated about 3.7 Mya (95% HPD: 5.5–2.2 Mya, Fig.2a).

The final nuDNA alignment consisted of 252 phased sequences (600 bp) for *mc1r* and 164 sequences (728 bp) for β*-fibint7* (accession numbers XX). A test for recombination using the Pairwise Homoplasy Index (*phi*) did not find significant evidence for recombination in either of the two genes (*p*-value = 0.772 for *mc1r*; *p*-value = 0.559 for β*-fibint7*). The nuDNA did not show a phylogenetic pattern fully congruent with the mtDNA, but a certain level of accordance can be detected for the β*-fibint7* fragments (Fig. 2c). In those specimens assigned to the newly detected mtDNA clade P, all but one (F41) β*-fibint7* allele form a monophyletic group. Moreover, all but three β*-fibint7* alleles (F6, F20 and F28) were found only in specimens belonging to the same mtDNA clade. The *mc1r* alleles were typically shared among specimens of different mtDNA lineages. However, most alleles identified in individuals of the western Pontine Islands are closely related at this locus and are mostly restricted to these islands, while they are very rare or absent elsewhere.

### Among island structure and gene flow

The *Podarcis siculus* populations of the western Pontine Islands were characterized by 31 mtDNA haplotypes with 44 polymorphic sites. The number of haplotypes (H) and nucleotide (π) and haplotype (h) diversity are reported for each dataset and marker in Table 1.

**Table 1.**
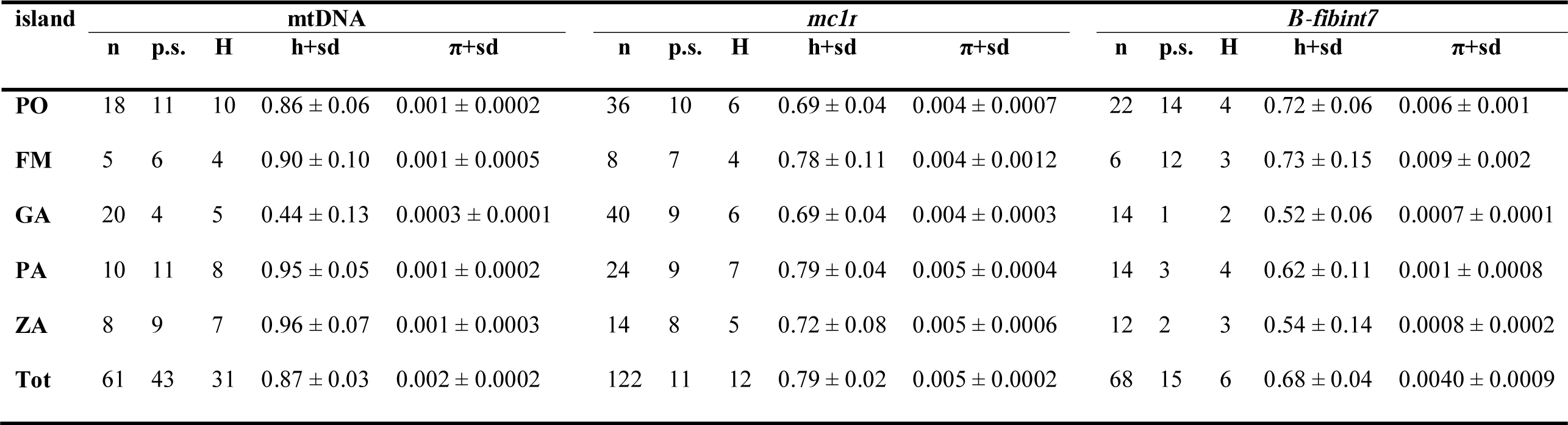
Genetic diversity estimates for each *Podarcis siculus* insular population (PO: Ponza, GA: Gavi, PA: Palmarola, ZA: Zannone). n; number of individuals/alleles, p.s.; number of polymorphic sites, H; number of haplotypes, h; haplotype diversity, π; nucleotide diversity, sd; standard deviation.

The mtDNA parsimony network showed the presence of two main genetic assemblages (Fig. 3a), one of which is restricted to the Zannone Island, where almost all specimens carried haplotypes of that assemblage. This result was also confirmed by the DAPC analysis, which suggested *K*=2 as the most reliable number of genetic clusters (Supplementary Fig. S2).

**Figure 3.**
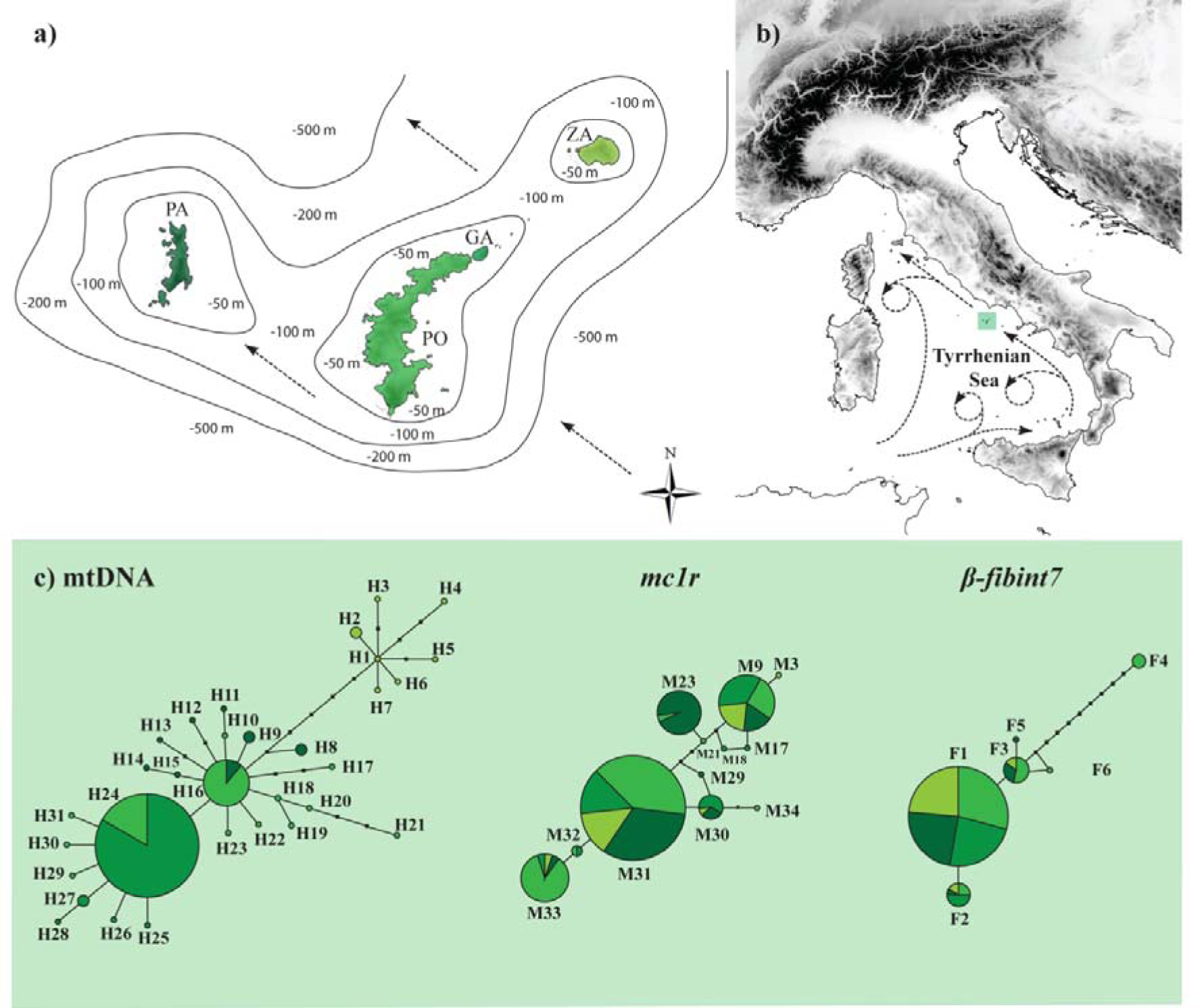
**(a)** Map and bathymetry of the western Pontine Islands (PO: Ponza, GA: Gavi, PA: Palmarola, ZA: Zannone). **(b)** Geographic location of the western Pontine Islands with the direction of the main marine currents in the Tyrrhenian Sea according to El-Geziry and Bryden^77^**. (c)** Statistical parsimony networks of the mtDNA (*cytb* + *nd4*) and nuDNA (*mc1r* and β*-fibint7*) fragments. Circles sizes are proportional to the haplotype frequencies and filled rectangles representing one mutation step. Haplotypes are colored by islands with different shades of green (for color code, see Fig. 3c).

The nuDNA parsimony networks did not identify a concordant pattern for either mitochondrial or island repartition (Fig. 3a). The *mc1r* fragment showed the presence of 12 alleles and most of them were found in individuals across all island populations. Notably, of the 6 β*-fibint7* alleles found, one allele (F4) was separated by 10 substitutions from the other 5 alleles. The DAPC carried out on nuclear sequences did not identify an optimal *K* value, returning a flat BIC curve (Supplementary Fig. S2).

All the 61 *P. siculus* specimens were genotyped at 11 polymorphic microsatellite loci, but the loci Pli 21 and Lv4a were excluded from further analyses because the former was monomorphic, while the latter might be influenced by positive selection. Genetic characteristics of the microsatellite loci are reported in Supplementary Table S6. Significant departures from the Hardy-Weinberg equilibrium (HWE) were observed in 2 out of 45 locus/population exact tests (Bonferroni correction applied: p<0.001, see Supplementary Table S7). No linkage disequilibrium was detected between any pair of loci. Evidence of null alleles was found for several loci, but none were consistent across all populations, therefore we did not exclude them from further analysis. An indication of recent reductions in population size (bottlenecks) was obtained only for the Faraglioni della Madonna population by both the Wilcoxon test (P = 0.00049) and the mode-shift indicator, which revealed a distortion in the typical L-shape distribution of allele frequencies.

Both clustering analyses conducted on microsatellite data (DAPC and STRUCTURE approaches) assigned *K*=2 as the most likely number of genetic clusters among islands (Supplementary Fig. S2 and Fig. S3). The first genetic cluster includes only the population from Gavi, while the second cluster includes all the remaining populations. Despite a further subdivision scheme, as suggested by the deltaK when *K*=4, the individual assignment at the fourth cluster did not correspond to any specific island, suggesting a reliable genetic partition not higher than two (Supplementary Fig. S3). The presence of further differentiation among all *P. siculus* populations is suggested by F_ST_ analyses, where all comparisons between populations yielded significant values (P<0.005) (Bonferroni correction applied; Table 2).

**Table 2.** F_ST_ values (below) and respective *p*-values (above) for each pair of populations (PO: Ponza, GA: Gavi, PA: Palmarola, ZA: Zannone). Significant differences after Bonferroni correction (p < 0.005) are emboldened.

|  | **ZA** | **GA** | **PO** | **FM** | **PA** |
| --- | --- | --- | --- | --- | --- |
| **ZA** | - | <0.00001 | 0.0048 | 0.0020 | 0.0010 |
| **GA** | **0.1144** | - | <0.00001 | <0.00001 | <0.00001 |
| **PO** | **0.0466** | **0.0655** | - | <0.00001 | 0.0010 |
| **FM** | **0.1859** | **0.1595** | **0.1081** | - | 0.0010 |
| **PA** | **0.0710** | **0.0807** | **0.0394** | **0.1347** | - |

The analysis of current gene flow performed with BayesAss indicated no evidence of recent gene flow in any pairwise comparisons. The migration rate estimates were all below 0.07 with the exception of the migration estimate between the Zannone and Palmarola populations. This was detected above the background noise (M = 0.16). The MIGRATE-N analysis carried out on the microsatellite dataset under a full migration matrix model indicated the presence of asymmetric gene flow among island pairs, with higher migration rates detected between the island of Ponza and all the others (Fig. 4). The Bayes factor calculated the model probability using the log marginal likelihood of each generated migration scenario and indicated that the longitudinal migration model was the most reliable scenario for gene exchange.

**Figure 4.**
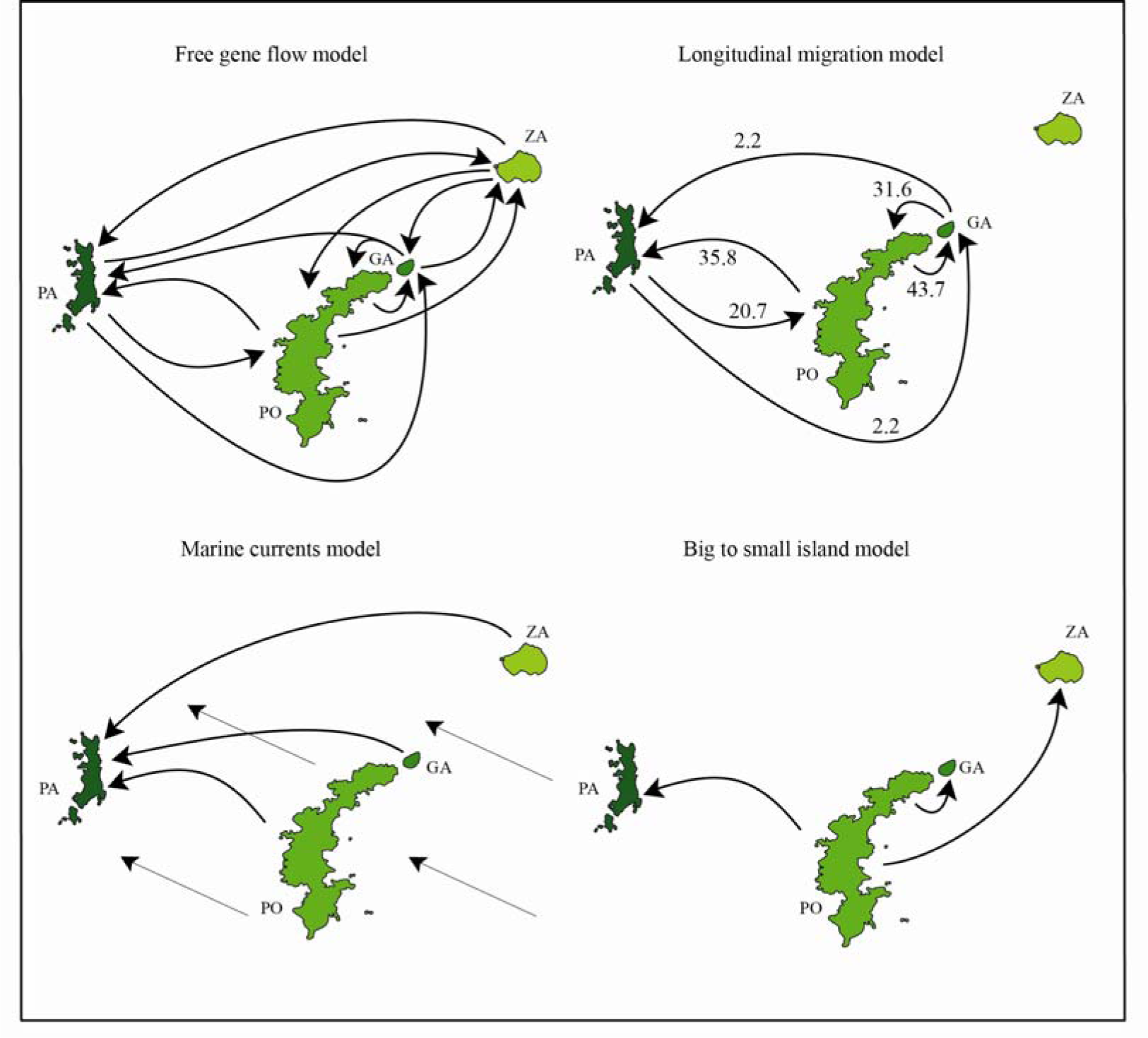
Models of migration among western Pontine Islands tested through Bayes factor analysis (PO: Ponza, GA: Gavi, PA: Palmarola, ZA: Zannone).Inferred migration rates are only shown for the most supported model.

## Discussion

Small island systems located on continental shelves have been influenced by cyclical sea level fluctuations and represent a compelling setting to investigate the birth and the formation of intraspecific genetic diversity within archipelagos. Although less explored compared to the attractive oceanic islands, continental and land-bridge archipelagos may offer comparable or even more dynamic systems^22^. For example, studies on Mediterranean islands underline a variety of microevolutionary processes, such as vicariance, dispersal, and secondary contact with gene flow, which may have contributed in parallel in assembling insular biota^9,11,15,24^. Our results fit with this emerging body of insular genetic diversity studies and outline an ancient dispersal event of *P. siculus* from the mainland to islands, followed by intra-islands vicariant processes with gene flow.

### On the origin of island genetic divergence

Phylogenetic reconstruction using two mitochondrial markers confirmed the complex phylogeographic pattern across the Italian Peninsula, with seven very divergent clades whose evolutionary histories have been described in detail in previous works^18,19^. In this study, we detect the presence of a new divergent 8^th^ *P. siculus* clade corresponding to populations inhabiting the western Pontine Islands of Ponza, Gavi, Palmarola, and Zannone. On the contrary, the island of Ventotene of the eastern Pontine Island group showed haplotypes belonging to the A2 clade, indicating a recent introduction of *P. siculus* to this island, as already reported^25^.

According to our estimate of divergence time, the ancestry of the Pontine clade dates back to the early/late Pliocene transition (3.7 Mya; 95% HPD: 5.5–2.2 Mya). Such a reconstruction is in line with the geological evolution of the western Pontine Islands (Fig. 2a). Indeed, on the basis of potassium-argon dating, the first volcanic event that accompanied the formation of the Ponza Island occurred between 4.2 and 3.7 Mya, followed by a quiescent period up to 3.2 Mya^21^. Although the estimated dates are associated with large uncertainty values of the coalescent process, our molecular clock suggests that the western Pontine populations colonized the paleo-archipalago early after their Pliocene volcanic formation. Furthermore, global changes in ice volume and records of terrestrial and marine sedimentary sequences suggest four glaciation events during the Pliocene, with one occurring about 3.6 Mya during the early/late Pliocene transition^26^. A consequence of global cooling is the sea-level retreat, which likely relaxed the separation of Ponza Island from the mainland facilitating a possible dispersal event. A similar explanation has also been forwarded to explain other ancient colonization events in the Mediterranean Basin^15,27^. It must be emphasized, though that the origin of these islands only puts an upper limit to the time since colonization and the age of the respective evolutionary lineages, as there is evidence that other vertebrate species colonized these islands much later^28^.

The genetic diversity of *P. siculus* from the Pontine Islands has been previously investigated using allozyme^29^. These authors were unable to detect any genetic differentiation of the western Pontine Island populations, neither with respect to the eastern Pontine Islands nor in comparison with the mainland populations. They subsequently suggested a possible human-mediated introduction in historical times. Our results from the nuDNA dataset clearly reject such a hypothesis, as allele frequencies of western Pontine Islands were significantly different from the other populations and several alleles of both *mc1r* and β*-fibint7* fragments were private either for the western Pontine Islands or even for single islands. The population structure defined by β*-fibint7* was more similar to the mtDNA, as all island alleles except one (F41) appear to form a monophyletic assemblage and no allele is shared with the mainland. On the contrary, most alleles of the *mc1r* gene occurred in specimens belonging to several mitochondrial clades. Such discrepancy is not surprising if we bear in mind that coding genes, such as *mc1r*, have lower mutation rates and are thus more prone to exhibit incomplete lineage sorting^30,31^. In addition, a recent study which reconstructed the phylogenetic relationship of 934 sequences of *mc1r* in nine *Podarcis* species revealed only slight differentiation, with some alleles shared even between different species^32^, corroborating our interpretation of incomplete lineage sorting.

### Inter-island genetic diversification

On the basis of historical sea level changes from the Early Pleistocene, the Pontine Islands have probably been subjected to repeated cycles of connection (into a single paleo-island setting) and isolation ^33^. Our inter-island diversification reconstruction is in line with such a scenario only if we consider a Middle Pleistocene divergence followed by different extent of genetic exchange during subsequent periods. The absence of a more ancient divergence is surprising considering the ancestry of these populations and the fragmentation of the islands assumed for the Middle Pleistocene periods due to sea level elevation. At the same time, it is also difficult to imagine that migration processes have been maintained for such an extended period (from 3.7 to 0.2 Mya) to prevent further divergence. A more plausible explanation is that populations diverged in allopatry, i.e., when islands were disconnected. However, some of these ancestral island populations may have become extinct during strong marine transgressions, such that their present population originates from a more recent divergences. Such an interpretation is also supported by stratigraphic and paleontological evidence, which suggest that at least two principal episodes of strong marine transgression on the Tyrrhenian Coast occurred some 0.54 and 0.35 Mya ago, which prompted a sea level rise from 55 up to 120 m a.s.l.^34^. In addition, beach deposits from the Early Pleistocene are present at an elevation of 200 – 250 m a.s.l. on Monte Guardia (Ponza Island), while abrasional surfaces from the Middle Pleistocene are widespread at 100 – 120 m a.s.l^35^. Therefore, considering the low altitude of some Pontine Islands (maximum altitude: Gavi = 60 m; Zannone = 170 m; Palmarola = 180 m; Ponza = 250 m), some of them might have been totally or partially submerged. Subsequent re-colonization from nearby islands during marine regressions may then have homogenized the genetic make-up among island populations.

According to our chronogram, the first episode of divergence within the Pontine Islands involved the separation of the Zannone Island population during the Mindel–Riss interglacial period (MIS 11, Fig. 2a). This period has been described as one of the longest warm periods of the Quaternary, inducing marked footprints of global sea level rise^36^. Under such vicariant mechanisms of divergence, taking into account similar distances and bathymetric depths among islands, we should expect also a similar extent of genetic diversification followed by genetic admixture as a consequence of glacial inter-island reunification. Indeed, sea depth between islands never exceeds 100 m (Fig. 3b), which is almost the same magnitude of sea level drop during the last glacial maximum^37,38^.

Our multilocus dataset revealed a pattern of mito-nuclear discordance within the western Pontine Islands. The mtDNA showed the presence of two principal assemblages: one composed of individuals from Zannone Island and the other including individuals from Ponza, Gavi, and Palmarola islands. However, the nuDNA did not support strong population structure, neither when relating it to geography (different islands) nor to the mtDNA groups. Finally, the clustering analysis carried out with both DAPC and STRUCTURE on the microsatellite dataset identified a *K*=2 subdivision, revealing the Gavi Islet as a separate cluster. The lack of accordance between genes and genealogies represents a common issue, but can provide relevant insights into microevolutionary processes underlying the origin of small island system biodiversity^39^. In such an insular context, the causes of mito-nuclear discordance should be primarily a result of the different evolutionary trajectories of distinct markers. Indeed, the mtDNA has an inherently higher mutation rate and, due to its uniparental inheritance, is expected to reach fixation fourfold faster than the nuDNA genome because of its smaller effective population size and the associated higher susceptibility to demographic changes and genetic drift. On the contrary, the lack of congruence of the nuDNA dataset to both geographic settings and mtDNA differentiation is probably because genetic differentiation on these islands is rather recent, such that the sorting of the nuclear genealogies is not complete. Such a scenario has been repeatedly reported in many vertebrate species, including other *Podarcis* lizards^30,31^.

Although a principal trend of mito-nuclear discordance emerged when considering the mere genealogies and the current structure of the markers, our coalescent-based analysis of gene flow allowed us to partially explain this complexity. Indeed, the most supported migration scenario, through Bayes factor calculation, was a longitudinal migration model. This model suggests reduced gene flow between the Ponza and Zannone islands, which are lengthwise arranged, and higher gene flow between the Ponza and Palmarola islands, which show a parallel arrangement that may have favored gene exchange (Fig. 3b). Such a prediction, further in line with the mtDNA signature, is particularly plausible because the genetic exchange could have been even more pervasive during a low sea level stand, when islands shared a larger contact area. However, we cannot rule out that the reduced gene flow between the Ponza and Zannone populations is the outcome of demographic dynamics like “density blocking”^40,41^, or alternatively, to morphological or behavioral changes which occurred during the allopatric divergence.

## Conclusion

The genetic differentiation we found in the populations from the western Pontine Islands with respect to the mainland is comparable with those observed for many other *Podarcis* lizards species and over twice as high as between the insular endemic *P. raffonei* and its sister taxon *P. wagleriana*^24,42,43^. On the other hand, populations from the eastern Pontine Islands seem to have colonized the Ventotene Island in recent times^25^. Considering the greater distance from the mainland and the deeper water in which the eastern Pontine Islands are located, the absence of a comparable ancient lineage is fairly surprising. Interestingly, in 1926 the herpetologist Robert Mertens described a particular phenotype endemic to Santo Stefano Island (*P. siculus sanctistephani*), which then became extinct and was replaced by the ordinary mainland phenotype during the first decades of the last century^44,45^. Such observations deserve special attention and may have several implications. First, it could be argued that an ancient sister lineage was historically present on the eastern Pontine Islands, including Ventotene. Secondly, and perhaps more fascinating, would be to understand why the *P. siculus sanctistephani* became extinct. Currently, there is no trace of introgression in the current allochthonous population from the Santo Stefano Island island^25,29^. If the local endemic *P. siculus sanctistephani* became extinct after the new colonizers and was indeed not hybridizing with later arriving conspecifics of the mainland type, there might have been a reproductive isolation mechanism, e.g. a behavioral or morphological change present in these island populations.

All these considerations point to a priority of the conservation of all the western Pontine Island populations, which should be referred to as *Podarcis siculus latastei* (Bedriaga, 1879). Indeed, the aforementioned example of possible population replacement underscore the vulnerability of endemic insular populations suggesting that further introductions from the mainland could be deleterious and should be prevented in order to maintain the genetic integrity on this archipelago. In support of our inferred genetic separation, morphological, behavioral, and physiological studies would be needed to unravel whether the found genetic divergence is associated with ecological and/or phenotypic diversification on these islands.

## Materials and methods

### Laboratory techniques and molecular data

A total of 156 specimens of *P. siculus* have been analyzed in this study: 63 new tail tissues were collected from six islands (Ponza, Faraglioni della Madonna, Gavi, Zannone, Palmarola, and Ventotene) while 93 individuals from 46 localities of the Italian Peninsula have been selected from a previous study^19^ (see Supplementary Table S1).

Genomic DNA was extracted according to the universal protocol of Salah^46^. We amplified one portion of the NADH dehydrogenase subunit 4 (*nd4*) with flanking regions (tRNA^Ser^, tRNA^His^ and tRNA^Leu^) in all 156 collected samples. Additionally, three fragments including the cytochrome b (*cytb*), the coding melanocortin receptor 1 (*mc1r*), and the β-fibrinogen intron 7 (β*-fibint7*) were amplified only in the 61 specimens of the Pontine Islands, as the samples from the Italian Peninsula had been previously analyzed at these loci by Senczuk et al. (2017). Details on reaction buffer, primers, and cycling conditions are reported in Supplementary Table S2. The obtained amplified products were purified with Sure Clean (Bioline) and sequenced with an automated DNA sequencer by Macrogen (www.macrogen.com).

BioEdit 7.2^47^ was used to evaluate the electropherograms, encode heterozygote positions using IUPAC ambiguity codes, and compute consensus alignments. The two mitochondrial fragments (mtDNA) were concatenated. For each nuclear gene (nuDNAs), we inferred the gametic phase using the coalescent-based Bayesian algorithm in PHASE 2.1^48,49^. The β*-fibint7* showed insertion/deletion (INDEL) polymorphisms in a region of poly-T and poly-G starting at 157 bp that, following the method described by Flot^50^, were used to determine the phase for sequences that were heterozygous for INDELs. The known phases were then implemented in the coalescent-based Bayesian reconstruction. Three independent runs were conducted with a burn-in at 1×10^3^, 1×10^4^ iterations, and thinning at each 100 steps. Finally, the occurrence of recombination events was assessed for each nuclear gene through the Pairwise Homoplasy Index (*phi*) test implemented in splitstree v. 4^51^.

The 61 samples from the western Pontine Islands were also genotyped at 11 microsatellite loci. Details on primer and PCR conditions are reported in Supplementary Table S3. Microsatellite multiplex amplifications were performed and then compared with singleplex PCR to confirm that there were no apparent allele dropouts or allele size differences. Amplification products were run on an ABI 3130xl Genetic Analyzer and allele size was scored using peak scanner 1.0 (Applied Biosystems). Amplification and genotyping were repeated twice for some 15% of the total samples (10 lizards) to verify reproducibility in microsatellite scoring. Genepop 3.4^52^ was used to assess deviations from Hardy-Weinberg equilibrium and linkage disequilibrium. The statistical significance levels were corrected for multiple comparisons using Bonferroni’s method (α =0.05). LOSITAN^53^ was used to identify candidate loci under selection. Before subsequent analyses, the presence of null alleles, large allele dropouts, and scoring errors was checked using MICRO-CHECKER^54^.

### Phylogenetic reconstruction and time of divergence

To infer phylogenetic relationships, the mtDNA dataset was analyzed using MrBayes v. 3.1.2^55^. The best substitution model for each partition (i.e., HKY + I+ G for *nd4* and GTR + I+ G for *cytb*) was selected using JModeltest^56^ under the Akaike Information Criterion (AIC). The analysis was run for 3×10^6^ generations using four annealing chains, sampling every 1×10^4^ and discarding the first 10% of computed trees as burn-in. Sequences of *Podarics muralis* retrieved from GenBank (accession numbers: HQ65293 for *cytb*, KF372413 for *nd4*) were used as an outgroup.

The software BEAST 1.8.1^57^ was used to date the most recent common ancestors (TMRCA). In this analysis the selection of proper priors is a crucial component although it may be challenging. Because of the absence of fossil records in *Podarcis* lizards to calibrate the phylogeny, we used substitution rates previously estimated for both gene fragments^12,43^. We incorporated a lognormal prior distribution to the mean rate in order to constrain the 95% of the prior distribution between the upper and lower values estimated for *Podarcis* (μ = 0.0175, SD = 0.2 for *cytb*, and μ = 0.0115, SD = 0.5 for *nd4*). Another relevant issue is the consideration of how substitution rates can vary across branches assuming a clock-like or alternatively a relaxed-clock model. To resolve this issue, a preliminary run was performed using a lognormal relaxed-clock in order to account for branch heterogeneity by screening the *ucld.stdev* parameter, which provides information about whether evolutionary rates inferred from the data are clock-like. Finally, the choice of an outgroup can be difficult when the phylogeny is not fully resolved^32^. However, the use of a relaxed-clock allows estimates of the tree root even in the absence of a known outgroup^58^. BEAST was run for 1×10^8^ generations sampling every 1×10^4^ steps and the software tracer 1.6^59^ was used to check parameter convergence. The final consensus tree was summarized from the stationary distribution (burn-in=25%) in TreeAnnotator 1.8.1^57^.

Furthermore, to better understand the relationship between mtDNA and nuDNA and the variation of the latter throughout the whole species distribution area, two nuDNA parsimony networks were built using the software TCS 1.21^60^ under the 95% probability connection limit. Finally, uncorrected pairwise *p*-distances were calculated for each mtDNA clade obtained by the phylogenetic reconstruction using MEGA 7.0^61^.

### Among island structure and gene flow

To infer the genetic relationship between mitochondrial and nuclear haplotypes among island populations, we used the mtDNA and nuDNA dataset to construct statistical parsimony networks under the 95% probability connection limit using the software TCS 1.21.

For the mtDNA and nuDNA datasets, the number of haplotypes (H) as well as nucleotide (π) and haplotype (h) diversity were estimated using DnaSP 5.1^62^. The microsatellite genetic variability was estimated by calculating allele frequencies, expected (H_E_) and observed (H_O_) heterozygosities, number of alleles, and number of private alleles using Genetix 4.05^63,64^ and Fstat 2.9.3.2^65^. All calculations were performed for the whole dataset and for each island. The software Bottleneck 1.2.02^66^ was used to assess the occurrence of recent bottlenecks; the tests were performed assuming the infinite allele model (IAM).

To explore the genetic structure among island populations, we performed a discriminant analysis of principal component (DAPC) implemented in the R-package ‘Adegenet’^67^ using the mtDNA, nuDNA, and microsatellite datasets. For the microsatellite data we also performed a Bayesian clustering analysis implemented in STRUCTURE 2.3.1^68^ to infer the number of different genetic clusters (*K*), assuming the admixture model. Ten simulations of 1×10^5^ iterations for each *K* (from *K*=1 to *K*=7) were run with a burn-in of 1×10^6^. The best *K* value, based on the modal value Δ*K*^69^, was detected using the online software Structure Harvester 0.6.93^70^. In addition, Arlequin 3.5^71^ was used to investigate the interpopulation level of genetic differentiation by calculating *F_ST_*, according to Weir and Cockerham^72^.

In order to assess the presence of gene flow and whether such transfer was recent or historical we employed two different approaches. The software BayesAss 3.0^73^ was used to estimate the presence of gene flow within the past few generations. Several preliminary runs were performed to adjust the acceptance rate of the mixing parameters (migration rates, allele frequencies, and inbreeding coefficients). The final analysis was carried out with 5 independent replicates running for 1×10^7^ iterations, sampling every 100 steps and discarding the first 1 million generations as burn-in. Convergence was assessed by comparing the accordance of the posterior mean parameter estimates across independent runs and by checking the stationary of the trace file using the software TRACER. To obtain an estimate of a more ancient signal of gene flow, we used the software MIGRATE-N^74,75^. The program implements a coalescent-based approach to estimate the mutation scaled effective population size (θ = 4N_e_*u*) and the mutation scaled long-term migration rate (M = *m/u*, where *m* is the migration rate, and *u* is the mutation rate). We generated a set of plausible a priori migration models, originated from insular biogeography, to test empirical data: (i) the “free gene flow model”, which implies gene exchange among all islands, (ii) gene flow among all islands except for Zannone, as could be expected between islands arranged in a longitudinal axis (“longitudinal migration model”), (iii) gene flow in a south-east/north-west direction as could be supposed when considering the marine currents, favoring overseas dispersal (“marine currents model”), and (iv) gene flow from the bigger island to the smaller (“big to small island model”). Bayes factors were calculated for each model and used to test the more reliable scenario following the guideline of Kass and Raftery^76^. The Faraglioni della Madonna population was excluded from this analysis because of the small sample size (n < 5). After a preliminary tuning run to estimate θ and *m* parameters through F_ST_ to use as starting values, we performed four independent runs for each generated model using four parallel chains with static heating (temperatures 1.0,1.5, 3.0 100,000.0) and Brownian motion microsatellite model.

## Data Availability

All DNA sequences used in this study will be submitted to GenBank upon acceptance.

## Acknowledgements

The authors are very grateful to Paolo Colangelo, Flavia Annesi, Ignazio Avella, and Laura Gramolini for their advice and laboratory support. Thanks are also extended to all the collectors who have contributed to improving our sampling. We wish to give a special thanks to Dario D’ Eustacchio, who tragically passed away, for his precious suggestions and for his valuable help in the field. We thank Rebecca Nagel for her “native-speaker’s” corrections of our manuscript. We are grateful to the Italian Environment Ministry for the Environment, Land and Sea for research permission (Prot. 00017879/PNM - 09/09/2012). Funding was provided by the Italian Ministry of Education, University and Research (PRIN 2012 project) and by “Progetti di Ricerca di Università” to RC, German Academic Exchange Service (DAAD) fellowship to GS, and by the Institute of Biochemistry and Biology, University of Potsdam to RT.

## Author contributions

G.S. and R.C. designed the study. R.C. and R.T. provided funding. G.S., R.C., and E.D. performed fieldwork and sample collection. G.S., K.H., C.R., and E.D. obtained the molecular data. G.S., V.M., and C.R. analyzed the data. R.T. guided and contributed to discussions on the phylogeographical and microsatellite analyses. G.S. wrote the manuscript. All authors reviewed and approved the manuscript.

## Additional information

### Supplementary information

**Table S1.** Locality names with their relative geographic coordinates, sample size, and frequencies of each mitochondrial and nuclear haplotype.

**Table S2** Primers used in the study with relative references and cycling conditions. PCRs were conducted in a standard volume of 25 µL, containing 1 mM Tris-HCL (pH 9.0), 5 mM KCL, 0.15 mM MgCl_2_, 0.2 mM of each dNTP, 0.1 mM of both forward and reverse primer, and 0.5 units of *Taq* polymerase.

**Table S3** PCRs were conducted in a standard volume of 25 μL, containing 1 mM Tris-HCL (pH 9.0), 5 mM KCL, 0.15 mM MgCl_2_, 0.2 mM of each dNTP, 0.1 mM of both forward and reverse primer, and 0.5 units of *Taq* polymerase. Number of asterisks indicate pair of loci amplifications.

**Table S4** Accession numbers of each gene fragment analysed in this study with relative haplotypes. In bold are sequences retrieved from Senczuk et al. 2017. * new sequences processed in this study (GeneBank accession numbers will be provided upon acceptance).
**Table S5.** Uncorrected pairwise *p-*distance calculated for each mtDNA clade obtained by the phylogenetic reconstruction.

**Table S6** Genetic characteristics of each microsatellite loci. N = number of individuals; R = size range of alleles in base pairs (bp); N_A_ = number of alleles, brackets indicate the number of private alleles; H_E_ = expected heterozygosity; H_O_ = observed heterozygosity; F_IS_ =
inbreeding coefficient; N_T_ = total number of alleles, brackets indicate the number of private alleles.

**Table S7.** P values for deviation from Hardy-Weinberg equilibrium. Bonferroni correction *p*<0.001.

**Figure S1.** Phylogenetic reconstruction of *Podarcis siculus* with MrBayes, based on concatenated data from *cytb* and *nd4.* The posterior probabilities of the main clades are shown at each node.

**Figure S2.** Values of BIC calculation versus number of clusters in the Discriminant Analysis of Principal Component (DAPC) for each analyzed dataset.

**Figure S3.** Evanno method in STRUCTURE HARVESTER (above) and the results of the Bayesian clustering analysis preformed in STRUCTURE based on 10 microsatellite loci for 5 insular populations of *Podarcis siculus* (below). Group repartition at *K*=2 and *K*=4 is shown according to the rate of change of the likelihood function (deltaK). The genetic clusters are coded by distinct colors and each vertical line represents the membership probability of one individual to a genetic cluster. On the horizontal bar, each sampled island is shown: PO = Ponza; PA = Palmarola; GA = Gavi; ZA = Zannone; FM = Faraglioni della Madonna.

